# Impact of N^6^-methyladenosine (m^6^A) machinery on HIV-1 replication in primary CD4+ T cells

**DOI:** 10.1101/2025.11.07.686373

**Authors:** Kathryn A. Jackson-Jones, Lacy M. Simons, Siyu Huang, Thea L. Joseph, Aubrey M. Sawyer, Li Wu, Judd F. Hultquist

## Abstract

N^6^-methyladenosine (m^6^A) is the most prevalent internal modification of cellular and viral RNA and is critical to the regulation of its localization, stability, and translation. Previous studies on the role of m^6^A during HIV-1 replication have produced conflicting results. Since m^6^A function can vary dramatically by cell type and state, here we aimed to clarify the role of the m^6^A machinery during HIV-1 replication in primary CD4+ T cells. Using CRISPR-Cas9 we targeted 46 cellular genes implicated in m^6^A or 5-methylcytosine (m^5^C) regulation and measured subsequent HIV-1 replication in primary CD4+ T cells. Only knockout of the m^6^A writer complex auxiliary proteins VIRMA and WTAP, and the m^6^A reader YTHDF2 were validated as significantly decreasing HIV-1 replication. In contrast, knockout of METTL3 or METTL14, which form the catalytic core of the writer complex, resulted in only marginal changes in HIV-1 infection, despite significant decreases in total cellular m^6^A levels. Chemical inhibition of METTL3 led to a dose-dependent decrease in HIV-1 infection, coupled with an increase in protein levels of METTL3 and other writer complex members. Expression of writer proteins was also co-dependent, revealing complex regulatory feedback mechanisms. Overall, these results clarify the role of epitranscriptomic machinery during HIV-1 replication in primary CD4+ T cells and suggest regulation by auxiliary members of the m^6^A writer complex is more influential than the function of the catalytic core itself on HIV-1 infection in primary CD4+ T cells.

**IMPORTANCE:** m^6^A is the most common chemical modification on cellular and viral RNA and regulates its stability, localization, and translation. m^6^A modification and its regulation varies dramatically between cell types and cell states. In this study, we investigated the role of m^6^A factors during HIV-1 infection of physiologically relevant primary CD4+ T cells. Using CRISPR-Cas9 to knockout 46 cellular genes implicated in RNA modification, we found only the m^6^A writer complex auxiliary members WTAP and VIRMA, and the reader YTHDF2, significantly affected HIV-1 replication in these cells. In contrast, knockout of METTL3 or METTL14, which form the catalytic core of the writer complex, resulted in marginal changes in HIV-1 infection, despite larger reductions in total cellular m^6^A levels. Our findings suggest regulation by auxiliary members of the m^6^A writer complex is more influential than the function of the catalytic core itself on HIV-1 infection in primary CD4+ T cells.

## INTRODUCTION

Post-transcriptional modification of cellular and viral RNA is critical to the regulation of its localization, stability, and translation (1–4). Over 300 distinct epitranscriptomic modifications have been described to date (5). These modifications are regulated by cellular proteins that can be broadly categorized as “writers” that facilitate the modification, “erasers” that remove the modification, or as “readers” that recognize the modification and direct downstream function (1–3,6,7). Collectively, these proteins and their associated regulatory complexes help determine the degree, location, and functional consequences of these modifications on their RNA targets.

The reversible conversion of adenosine to N^6^-methyladenosine (m^6^A) is the most prevalent co-transcriptional RNA modification in eukaryotic cells (1,8,9). The large m^6^A writer complex contains a heterodimer core of the catalytically active methyltransferase 3 (METTL3) bound to the stabilizing methyltransferase 14 (METTL14) (10–13). However, the heterodimer core alone has limited activity (14,15) and several other auxiliary writer complex members have been identified as necessary for stabilizing the writer complex and required for m^6^A modification (16,17). Binding of Wilms’ tumor 1 associating protein (WTAP) to the heterodimer regulates the methyltransferase activity or the core heterodimer (10,17) and has been shown to be essential for nuclear speckle localization in HeLa cells (18), suggesting that WTAP may recruit METTL3-METTL14 to their target mRNAs. Further members include Vir like m^6^A methyltransferase associated (VIRMA, also known as KIAA1429) (19–22), Zinc finger CCCH-type containing 13 (ZC3H13) (20,22,23), RNA binding motif protein 15 (RBM15) (20–22,24,25) and Casitas B-lineage lymphoma-like 1 (CBLL1, also known as HAKAI) (20–22,26,27). All have been shown to be required for m^6^A modification in various systems and suggested to direct target and positional specificity of the core heterodimer (19), however we have limited understanding of the specific roles played by each non-catalytic subunit. In contrast to the METTL3-METTL14 heterodimer, METTL16 is only able to methylate a limited number of RNAs (28–30) and METTL4 mediates internal m^6^Am methylation of small nuclear RNA (snRNA) (31).

The impact of RNA modification is heavily dependent on the network of m^6^A regulators in each cell type (32–34). As such, whilst m^6^A, and additionally 5-methylcytosine (m^5^C), modifications have been found to be important for human immunodeficiency virus type 1 (HIV-1) replication, reports often disagree on the extent, location, and functional importance of these modifications in a manner that is dependent on cell model and experiential approaches. For example, a study of HIV replication in the CEM-SS T cell line found that most m^6^A modifications were localized in the 3’ untranslated region (UTR) and that these modifications were critical for enhancing viral mRNA expression and thereby HIV-1 replication potential through the recruitment of the three YTH domain family (YTHDF) m^6^A readers; YTHDF1, YTHDF2 and YTHDF3 (35). In the MT4 T cell line, on the other hand, m^6^A modifications were reported to occur throughout the entire HIV genome with modifications in the Rev-response element (RRE) important for mediating viral RNA nuclear export through Rev, and a dramatic increase of host m^6^A levels upon infection (36). In the Jurkat T cell line, m^6^A modifications were found to occur primarily in the 5’ and 3’ UTRs of the HIV-1 genome and be bound by YTHDF readers, but only YTHDF1 (not the other YTHDF proteins) inhibited reverse transcription and only YTHDF3 (not the other YTHDF proteins) repressed viral transcription. Furthermore, in Jurkat and primary CD4+ T cells, HIV-1 infection did not significantly affect the total level of m^6^A on host RNA (37). However in CD4+ SupT1 T cells, HIV infection modified both the m^6^A and m^5^C host epitranscriptomic with a bias towards increased methylation (38). In HEK293T cells, m^6^A modifications were likewise found to be enriched at the 5’ and 3’ UTRs, with modification of the 5’UTR driving enhanced translation but interfering with viral genome packaging (39). In another report, m^6^A modifications were enriched in the 3’UTR of the viral genome in HEK293T cells, and bound by the reader YTHDC1 which regulated the alternative splicing of HIV-1 RNAs, and by the reader YTHDF2 which enhanced viral gene expression and stabilized viral RNAs in CEM-SS T cells (40). Finally, in differentiated human monocytic U937 cells and primary monocyte-derived macrophages, m^6^A modifications were found to shield viral transcripts from innate immune sensing and induction of antiviral cytokine type-I interferon (IFN) (41). Multiple functions have likewise been ascribed to m^5^C modification of the HIV viral genome, including promoting mRNA translation in CEM cells (42) and inhibition of Tat-dependent transcription in HeLa cells (43). It has even been suggested that the m^5^C writer NOP2/Sun RNA methyltransferase 2 (NSUN2) and the METTL3-METTL14 heterodimer may work cooperatively to enhance translation of specific RNAs (44). While several of these results and mechanisms are not mutually exclusive, these discrepancies highlight the need to better understand the role of these epitranscriptomic modifications directly in primary human cells.

In this study, we performed a systematic investigation of m^6^A and m^5^C machinery during HIV-1 replication in primary CD4+ T cells. First, we confirmed that the core m^6^A machinery was ubiquitously expressed in resting, activated, and IFN-stimulated primary CD4+ T cells and found that the eraser FTO is upregulated in response to T cell activation. We then used CRISPR-Cas9 ribonucleoproteins to knockout 46 genes previously implicated in m^6^A or m^5^C modification in primary CD4+ T cells from multiple donors, subsequently challenging the cells with replication-competent HIV-1. Of the screen hits, only knockout of the m^6^A writer complex auxiliary members VIRMA and WTAP, and the m^6^A reader YTHDF2 were validated as significantly decreasing HIV-1 replication with independent gRNAs and donors, suggesting they act as viral dependency factors. In contrast, knockout of METTL3 or METTL14, which form the catalytic core of the writer complex, resulted in marginal changes in HIV-1 infection, despite larger overall impacts on total cellular m^6^A levels, suggesting the auxiliary writer complex members may be aiding HIV-1 replication through a distinct mechanism. Overall, these results clarify the role of epitranscriptomic machinery during HIV-1 replication of primary CD4+ T cells and suggest regulation by auxiliary members of the m^6^A writer complex is more influential than the function of the catalytic core itself on HIV-1 infection in primary CD4+ T cells.

## MATERIALS & METHODS

### Cell lines and cell culture

Human embryonic kidney (HEK) 293T cells (ATCC, CRL-3216) were maintained in Dulbecco’s modified Eagle’s medium (Sigma-Aldrich, #D6429) supplemented with 10% heat-inactivated fetal bovine serum (FBS) (Gibco, #10082147) and 1% penicillin-streptomycin (50mg/ml; Corning). Cells were cultured at 37°C in a humidified incubator with 5% CO_2_.

### CD4^+^ T cell isolation and activation

Primary human CD4^+^ T cells from HIV-1 seronegative donors were isolated from either buffy coat (Vitalant) or leukopaks (STEMCELL Technologies). PBMCs were isolated by Ficoll centrifugation. Bulk CD4^+^ T cells were subsequently isolated from PBMCs by magnetic negative selection using an EasySep Human CD4^+^ T cell isolation kit (STEMCELL Technologies; #17952) according to the manufacturer’s instructions. Isolated CD4^+^ T cells were suspended in RPMI 1640 (Sigma-Aldrich) supplemented with 5mM HEPES (Corning), penicillin-streptomycin (50mg/ml; Corning), 5mM sodium pyruvate (Corning), and 10% FBS (Gibco), termed “complete RPMI”. Media were supplemented with interleukin-2 (IL-2; 20IU/ml; Miltenyi) immediately before use. For activation, bulk CD4^+^ T cells were immediately plated on anti-CD3-coated plates [coated for 12 hours at 4°C with anti-CD3 (20mg/ml) (UCHT1; Tonbo Biosciences)] in the presence of soluble anti-CD28 (5mg/ml; CD28.2; Tonbo Biosciences) and incubated for 72 hours at 37°C and 5% CO_2_. For RT-qPCR experiments, “untreated” cells were not exposed to CD3 and CD28 antibodies after isolation and were instead cultured in complete RPMI without IL-2, and “interferon (IFN)-stimulated” cells were stimulated with 1000 U/mL universal type I interferon (IFN) (PBL Assay Science, #11200-2) for 24 hours, post activation with CD3 and CD28 antibodies.s

### RNA extraction and reverse transcription quantitative polymerase chain reaction (RT-qPCR)

Total RNA was extracted from primary CD4+ T cells using QiaShredder columns (Qiagen, #79656) followed by RNeasy Mini kit (Qiagen, #74106) according to the manufacturer’s instructions. cDNA was synthesized from extracted RNA using Transcriptor Reverse Transcriptase (Roche #03531295001), following the manufacturer’s protocol. Following cDNA synthesis, transcript abundance was assessed by RT-qPCR. Each gene was measured using the relevant standard PrimeTime RT-qPCR Assay (IDT, **Table S1**). For expression level normalization, *TBP* was measured (IDT, Hs.PT.58v.39858774). The reaction was performed using PrimeTime Gene Expression Master Mix (IDT, # 1055770). The PCR cycles were as follows: 95°C for 3min followed by 40 cycles of 95°C for 5s and 60°C for 30s. Reactions were performed in technical triplicate. All primer sequences can be found in **Table S1**.

### CRISPR-Cas9 RNP production

CRISPR-Cas9 ribonuclear protein (crRNP) production and primary CD4^+^ T cell editing were performed as previously published (45). Briefly, gene specific lyophilized Edit-R predesigned CRISPR RNA (crRNA) (Dharmacon, **Table S1**) and Edit-R transactivating CRISPR RNA (tracrRNA) (Dharmacon, #U-002005-50) were each suspended at a concentration of 160μM in 10mM Tris-HCl (7.4 pH) with 150mM KCl. crRNA was mixed with tracrRNA at a 1:1 ratio and incubated for 30 min at 37°C to anneal. The crRNA-tracrRNA complexes were then mixed gently with 40μM Cas9 (UC-Berkeley Macrolab) at a 1:1 ratio to form CRISPR-Cas9 ribonuclear proteins (crRNPs). Aliquots of crRNPs were frozen in Lo-Bind 96-well V-bottom plates (E&K Scientific) at −80°C until use. For multiplex crRNP synthesis, multiple crRNA targeting the same gene were mixed in equal proportions before proceeding with tracrRNA addition as described above. All crRNAs used in this study were purchased from Dharmacon or derived from the Dharmacon predesigned Edit-R library for gene knockout (**Table S1**).

### Electroporation of primary CD4^+^ T cell cultures

Each electroporation reaction mixture consisted of 5×10^5^ activated T cells, 3.5μL crRNPs, and 20μL P3 electroporation buffer (P3 Primary Cell 4D-Nucleofector® X Kit S, Lonza, #V4XP-3032). After three days of activation, cells were suspended in fresh complete RPMI and counted. crRNPs were thawed and allowed to warm to room temperature. Immediately prior to electroporation, cells were centrifuged at 400×*g* for 5 min, supernatant was removed by aspiration, and the pellet was resuspended in 20μL of room-temperature P3 electroporation buffer (P3 Primary Cell 4D-Nucleofector® X Kit S, Lonza, #V4XP-3032) per reaction. 20μL of cell suspension was gently mixed with each crRNP and aliquoted into a 96-well electroporation cuvette. Cells were electroporated using a 4D-Nucleofector® Core Unit (Lonza, #AAF-1003B) with an attached 4D-Nucleofector^®^ 96-well Unit (Lonza, #: AAF-1003S) using pulse code EH-115. Immediately after electroporation, 100μL of prewarmed complete RPMI supplemented with IL-2 (20IU/ml; Miltenyi) was added to each well, and cells were allowed to recover for at least 30min in a 37°C cell culture incubator. Cells were subsequently moved to 96-well flat-bottomed culture plates prefilled with 100μL warm complete RPMI supplemented with IL-2 (20IU/ml; Miltenyi) and anti-CD3–anti-CD2–anti-CD28 beads (T-cell activation and stimulation kit; Miltenyi, #130-091-441) at a 1:1 bead-to-cell ratio. Cells were cultured at 37°C and 5% CO_2_ in a humidified cell culture incubator for four days to allow gene knockout and protein clearance, with additional complete RPMI supplemented with IL-2 (20IU/ml; Miltenyi) added after 48 hours. Cells from each treatment were replica-plated for harvest of protein lysates and infection in triplicate, To collect protein lysates for determination of knockout efficiency, 1×10^5^ cells from each polyclonal cell culture were pelleted, supernatant removed, and pellets resuspended in 50μL 2.5X Laemmli sample buffer [1.9 mL 0.5 M Tris-HCl pH 6.8 (Fisher Bioreagents, #BP153-1), 6 mL 50% glycerol (Fisher Bioreagents, #BP229-1), 3 mL 10% SDS (Corning, #46-040-CI), 250 µL β-mercaptoethanol (Fisher Chemical #O3666I-100), 50 µL 1% bromophenol blue (Fisher Bioreagents, #BP115-25), 18.8 mL Dulbecco’s Phosphate-Buffered Saline (DPBS) (Corning, #21-031-CV)]. Protein lysates were heated to 98°C for 20min before storage at −20°C until immunoblotting.

### Immunoblotting of primary CD4^+^ T cell cultures

Samples were thawed and 15μL of each was loaded onto 4-20% Criterion TGX SDS-PAGE protein gels (Bio-Rad) plus 5μL of PageRuler Plus prestained protein ladder (Thermo Scientific #26619). Gels were run at 150V for 90min until the ladder was sufficiently separated. Proteins were transferred to PVDF membranes by methanol-based electrotransfer (Bio-Rad Criterion blotter) at 90V for 2h. Membranes were blocked in 5% milk in PBS with 0.1% Tween 20 for 1h at room temperature prior to overnight incubation with primary antibody against the protein of interest or GAPDH as a protein loading control. All primer sequences can be found in **Table S1**. Goat anti-rabbit immunoglobulin (IgG) (Jackson Immuno, #111-035-003) or goat anti-mouse IgG (Jackson Immuno, #115-035-003) HRP-conjugated secondary antibodies were detected using Immobilon Western Chemiluminescent HRP Substrate (Millipore, #WBKLS0500) and imaged on Hyblot ES Autoradiography Film (Denville Scientific), or by chemiluminescence using iBright FL1500 Imaging System (ThermoFisher). Blots were incubated in 1X ReBlot Plus Antibody Stripping Solution (Millipore) before reprobing.

### Preparation of HIV-1 stocks

Replication-competent reporter virus stocks were generated from an HIV-1 NL4-3 molecular clone in which GFP had been cloned behind an IRES cassette following the viral nef gene (NIH AIDS Reagent Program, #11349). Briefly, 10µg of the molecular clone plasmid was transfected (PolyJet; SignaGen) into 5×10^6^ HEK293T cells (ATCC, CRL-3216) according to the manufacturer’s protocol. 25mL of supernatant was collected at 48 and 72 hours and combined. The virus-containing supernatant was filtered through 0.45-mm polyvinylidene difluoride filters (Millipore) and precipitated in 50% polyethylene glycol [average molecular weight 6000; Sigma-Aldrich] and 0.3M NaCl for at least 6h at 4°C. Supernatants were centrifuged at 3500rpm for 20 min, and the virus was resuspended in 50μL of DPBS (Corning, #21-031-CV). Aliquots were stored at −80°C until use.

### HIV-1 p24 ELISA

Viral stocks were heat inactivated at 60°C for 30 minutes, then quantified using an HIV-1 Gag p24 Quantikine ELISA kit (R&D Systems, #DHP240B) according to manufacturer protocol. Samples were run in duplicate at dilutions of 1:500 and 1:2500. Viral stock concentrations were determined using a standard curve.

### HIV-1 infection of primary CD4^+^ T cell cultures

Four days post-electroporation or 24 hours post drug treatment, activated primary CD4^+^ T cells were plated into a 96-well, round-bottom plate at a cell density of 1×10^5^ cells per well in 200μL of complete RPMI as described above, supplemented with IL-2 (20 IU/ml). For high throughput screening assays in **Figure 2&3**, 2.5μL of concentrated virus stock was added to each well, diluted in complete RPMI. For all other experiments, 0.2ng of p24 quantified virus was added to each well, diluted in complete RPMI. At two, five and seven days post-infection (dpi), 75μL of each culture was removed, mixed 1:1 with freshly made 2% formaldehyde in PBS (Sigma-Aldrich) to fix and stored at 4°C until measurement by flow cytometry. Cultures were then supplemented with 75mL of complete IL-2–containing RPMI 1640 medium and returned to the incubator.

### Flow cytometry and data analysis

Flow cytometry analysis was performed on an Attune NxT Flow Cytometer (Thermo Fisher Scientific), recording all events in a 50μL sample volume after one 150μL mixing cycle. Data were exported as FCS3.0 files using Attune NxT Software v3.2.0 and analyzed with a consistent template on FlowJo™ Software (BD Biosciences) (**Figure S1**). For viability data, cells were gated to exclude cell debris by light scatter and the number of cells recorded for each condition. For analysis of infection data, cells were gated for lymphocytes by light scatter followed by doublet discrimination in both side and forward scatter. Cells with equal fluorescence in the BL-1 (GFP) channel and the VL-1 channel were identified as autofluorescent and excluded from the analysis. A consistent gate was then used to quantify the fraction of remaining cells that expressed GFP.

### m^6^A ELISA

Primary CD4+ T cells from three independent donors were edited with crRNPs targeting METTL3, METTL14, VIRMA, WTAP, YTHDF2 or non-targeting (NT) control, or treated with 0.5µM of the METTL3 inhibitor STM3006 for 24 hours. All cells were subsequently infected with replication competent NL4.3 HIV-1 Nef:IRES:GFP at 0.4ng p24 equivalent virus per 100,000 cells. Infection was monitored by immunostaining and flow cytometry as above and achieved between 2-5% infection in each donor. Four dpi, total RNA was extracted from 6×10^6^ primary CD4+ T cells using a RNeasy Mini kit (Qiagen, #74106). mRNA was isolated using Oligo (dT)_25_ bead selection (Dynabeads® mRNA Purification Kit, # 61006, ThermoFisher Scientific) according to manufacturer’s protocol. Global m^6^A levels were quantified in 50ng mRNA using a m^6^A RNA methylation ELISA protocol as previously described (34,46,47).

### Quantification and statistical analysis

Data were analyzed using GraphPad Prism 10 (GraphPad Software, La Jolla, CA). Error bars represent mean ± standard deviation unless otherwise stated. For all figures: * = p<0.05, ** = p<0.01, *** = p<0.001, **** = p<0.0001. Significance reported for differences in mRNA expression measured by RT-qPCR were calculated using an ordinary one-way ANOVA with Dunnett’s post-hoc multiple comparisons test comparing each treatment to the relevant untreated sample for each donor, with three technical replicates and two independent, biological replicates (**Figure 1C**). Western blot images were quantified with FIJI (Fiji is just ImageJ)(48) using the Analyze Gels tool. Each band was normalized to the GAPDH loading control for that lane and then compared to the NT condition (**Figures 2B & 4C**). The threshold for hits reported for differences in HIV-1 infection levels from the CRISPR-Cas9 knockout arrayed screen were calculated as those more than log2(1) fold change from the median at each time point, with three technical replicates and four independent, biological replicates (**Figure 2E**). Significance reported for differences in HIV-1 infection levels from follow up CRISPR-Cas9 knockout validation were calculated using an unpaired, two-tailed t-test comparing each individual knockout to the NT control, followed by Bonferroni correction for multiple comparisons, with three technical replicates and four independent, biological replicates (**Figures 3C**). Significance reported for differences in HIV-1 infection levels upon treatment with a range of STM3006 concentrations were calculated using an ordinary one-way ANOVA with Dunnett’s post-hoc multiple comparisons test comparing each treatment to DMSO treatment, with three technical replicates and three independent, biological replicates (**Figures 4A & 4B**). Significance reported for differences in HIV-1 infection levels upon CRISPR-Cas9 knockout were calculated using an ordinary one-way ANOVA with Tukey’s post-hoc multiple comparisons test comparing all treatments, with three technical replicates and three independent, biological replicates (**Figures 4C & 4D**). Significance reported for differences in global m^6^A levels were calculated using an ordinary one-way ANOVA with Dunnett’s post-hoc multiple comparisons test comparing each treatment to NT, with three independent, biological replicates (**Figure 4F**).

**Figure 1.**
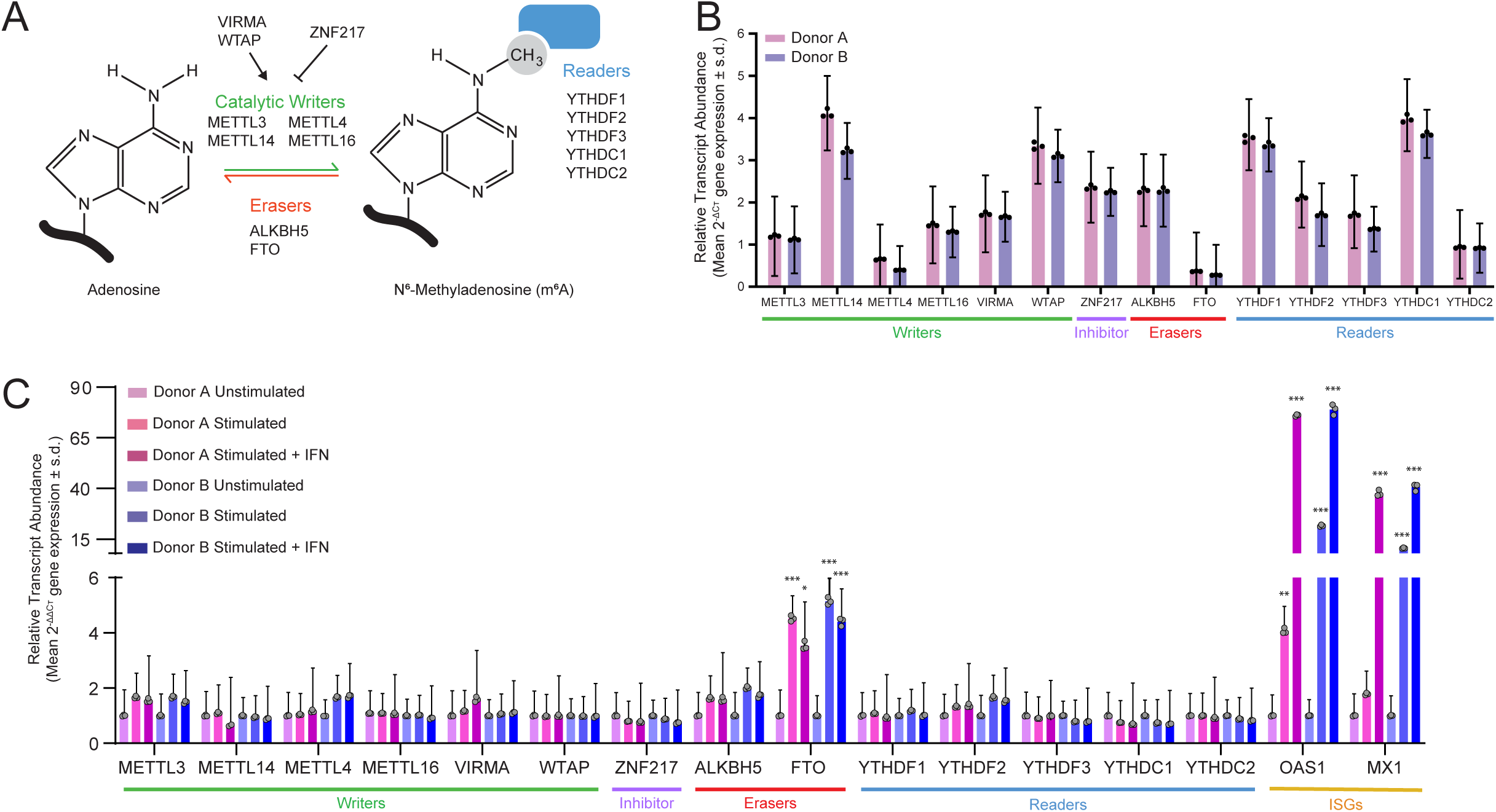
mRNA of the core m^6^A regulatory genes are expressed in primary CD4+ T cells. **A)** Schematic of the core m^6^A regulatory machinery. **B)** Relative transcript abundance of the indicated m^6^A regulatory genes relative to the housekeeping gene *TBP* (ΔCT) measured by reverse transcription quantitative polymerase chain reaction (RT-qPCR) in untreated primary CD4+ T cells isolated from two independent, HIV-1 seronegative donors. Bars show 2^-of mean cycle threshold (Ct) values normalized to the *TBP* housekeeping gene (ΔCT). Each dot represents one technical replicate. Error bars show the ΔCT standard deviation of technical triplicates. **C)** Relative transcript abundance of the indicated m^6^A regulatory genes, and two interferon (IFN)-stimulated genes, relative to the housekeeping gene *TBP* (ΔCT) measured by RT-qPCR in untreated, activated, and activated plus type I IFN-stimulated primary CD4+ T cells from two independent donors, normalized to untreated expression levels (ΔΔCT). Data represent the mean +/- ΔCT standard deviation of technical triplicates. Each dot represents one technical replicate. Error bars show the ΔCT standard deviation of technical triplicates. Statistics were calculated by Ordinary one-way ANOVA with Dunnett’s multiple comparisons test post-hoc, comparing each treatment to the relevant untreated sample for each donor.

**Figure 2.**
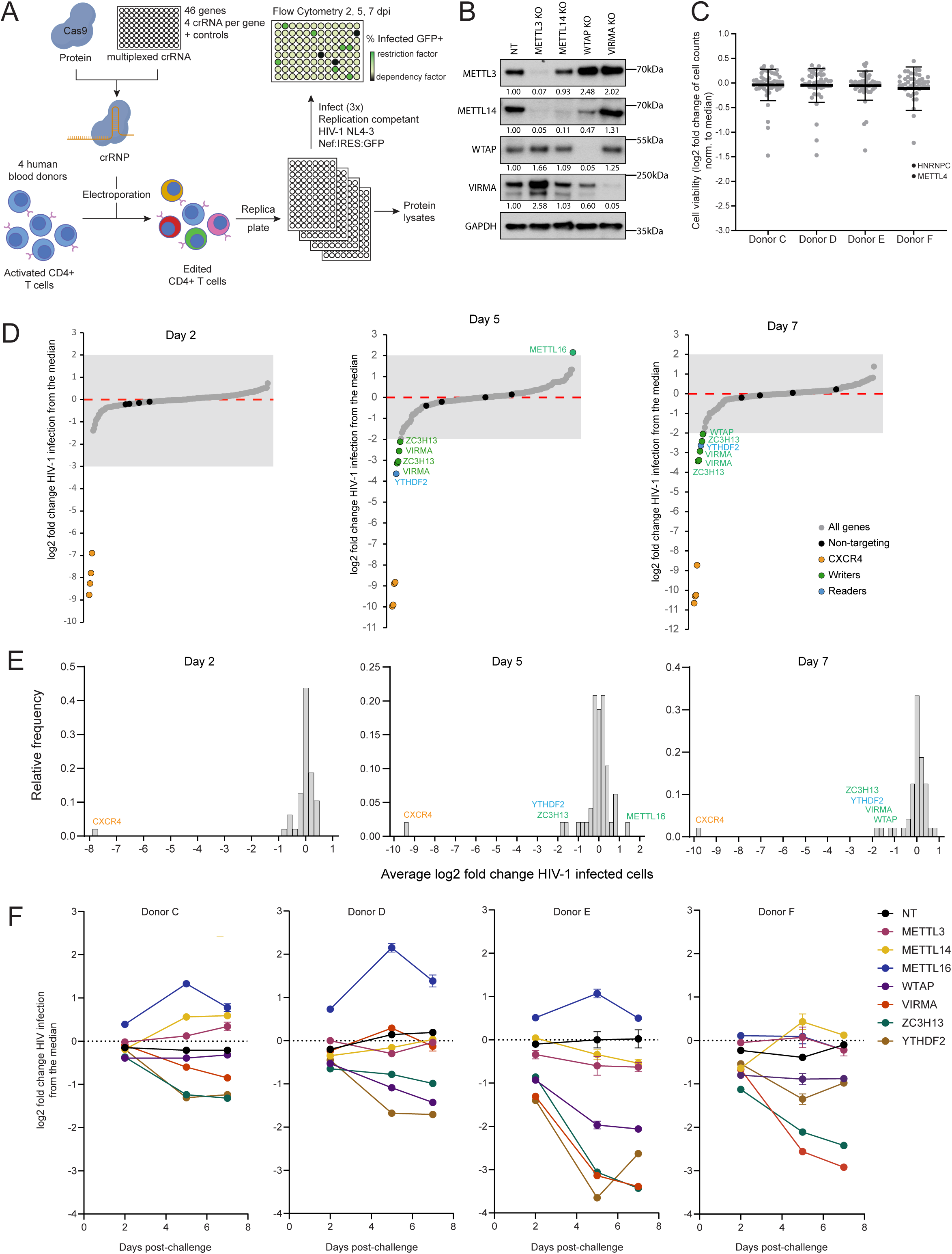
CRISPR-Cas9 knockout screen in primary CD4+ T cells identifies m^6^A regulators involved in the HIV-1 lifecycle. **A)** Schematic of the multiplexed CRISPR-Cas9 gene editing approach used on activated, primary CD4+ T cells from multiple, independent, HIV-1 seronegative donors. Four CRISPR RNA (crRNA) per gene were multiplexed and mixed with transactivating CRISPR RNA (tracrRNA) and Cas9 to form CRISPR-Cas9 ribonuclear proteins (crRNPs). Electroporation of the cells results in a polyclonal population of edited cells, which are replica plated for downstream assays. Four days post-electroporation, cells were infected in technical triplicate with replication-competent fluorescent reporter virus HIV-1 NL4-3 Nef:IRES:GFP and assessed by flow cytometry at two, five and seven days post-infection (dpi). **B)** Representative immunoblot of primary CD4+ T cells from one donor, demonstrating knockout efficiency of indicated m^6^A regulatory protein targets at four days post-electroporation with the relevant multiplexed crRNPs. Values below each band show the quantification, normalized to the GAPDH loading control and the NT condition. **C)** Cell viability of primary CD4+ T cells, shown as the log2 fold change in live cell number per knockout condition at six days post-electroporation relative to the plate median in four independent, HIV-1 seronegative donors. Each dot represents one knockout condition, line shows mean of all knockouts and error bars show standard deviation. Outliers (>1.5 log2 fold change from the median) are indicated as labeled black points and were excluded from downstream analysis. **D)** S-curve plots of the log2fold change in percent infected cells relative to the plate median for each knockout condition rank-ordered across all donors for each timepoint. Each dot represents one knockout condition. The dashed red line indicates the median; orange dots represent the *CXCR4*-targeting positive controls and black dots represent the non-targeting (NT) negative controls. Gene knockout with a log2 fold change greater than 2 are labeled and colored by category. **E)** Frequency histograms of log2 fold change in percent infected cells relative to the plate median for each knockout averaged across four donors at each timepoint. The threshold for hits reported was greater than a log2(1) fold change from the median at each time point. Knockouts meeting this criterion are labeled. **F)** HIV-1 infection over time in primary CD4+ T cells from four independent, HIV-1 seronegative donors after CRISPR-Cas9 knockout of the indicated subset of genes. Each dot represents average log2 fold change in HIV-1 infection from the median of three technical replicates and error bars indicate standard deviation.

**Figure 3.**
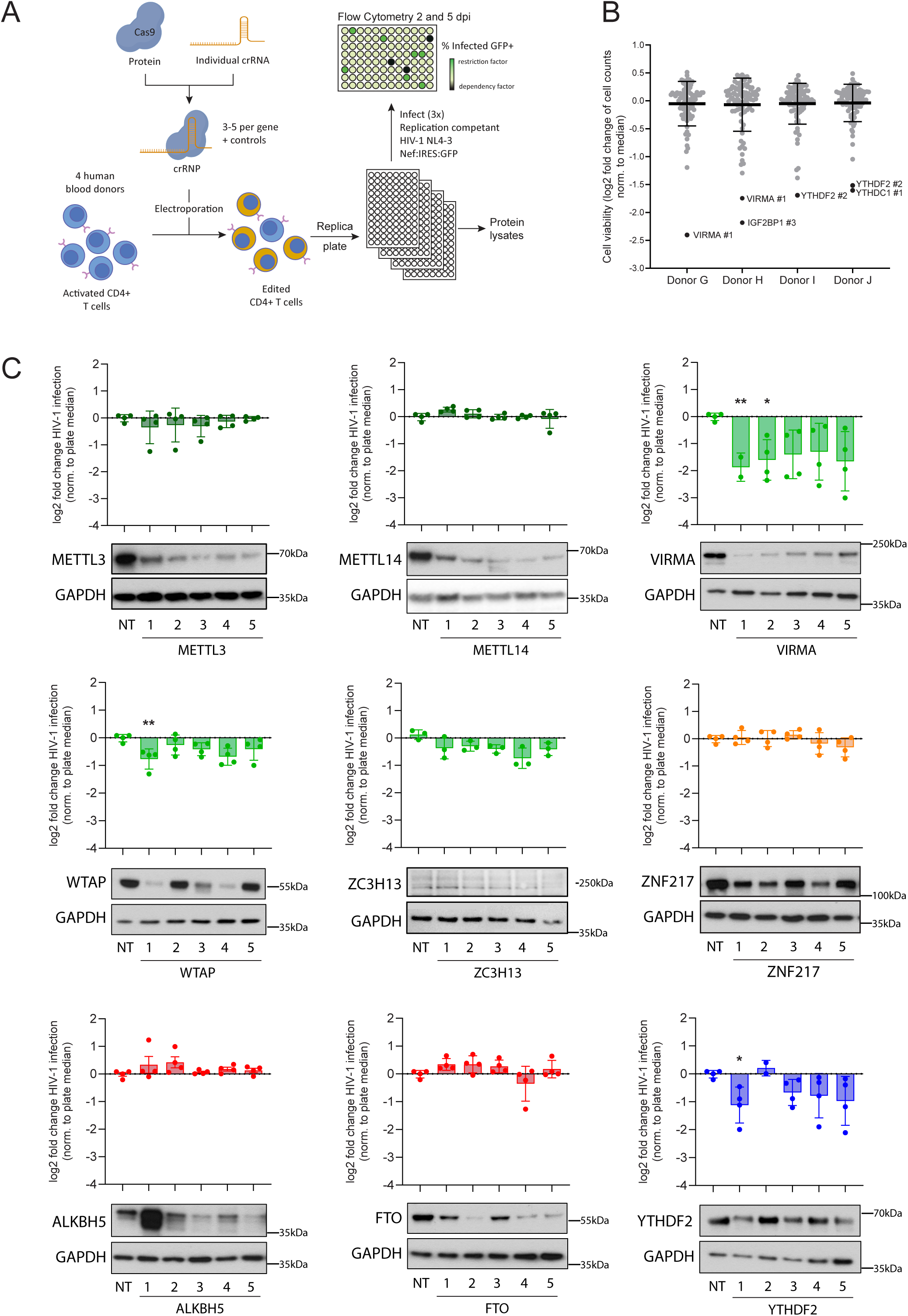
VIRMA, WTAP, and YTHDF2 validate as HIV-1 dependency factors. **A)** Schematic of the arrayed CRISPR-Cas9 gene editing approach used or phenotype validation in activated, primary CD4+ T cells from multiple, independent, HIV-1 seronegative donors. Between three and five independent CRISPR RNA (crRNA) per gene were separately mixed with transactivating CRISPR RNA (tracrRNA) and Cas9 to form CRISPR-Cas9 ribonuclear proteins (crRNPs), which are delivered in parallel. Electroporation of the cells results in a polyclonal population of edited cells, which are replica plated for downstream assays. Four days post-electroporation, cells were infected in technical triplicate with replication-competent fluorescent reporter virus HIV-1 NL4-3 Nef:IRES:GFP and assessed by flow cytometry at two, five and seven days post-infection (dpi). **B)** Cell viability of primary CD4+ T cells, shown as the log2 fold change in live cell number per knockout condition at six days post-electroporation relative to the plate median in four independent, HIV-1 seronegative donors. Each dot represents one knockout condition, line shows mean of all knockouts and error bars show standard deviation. Outliers (>1.5 log2 fold change from the median) are indicated as labeled black points and were excluded from downstream analysis. **C)** HIV-1 infection levels in primary CD4+ T cells five dpi with replication competent HIV-1, following knockout of the indicated genes. Data are displayed as log2 fold change in HIV-1 infection normalized to the plate median. Each dot represents the average infection from three technical replicates in one donor. Error bars show standard error of the mean. Significance levels were calculated using an unpaired, two-tailed t-test comparing each individual knockout to the non-targeting (NT) control, followed by Bonferroni correction for multiple comparisons. Representative immunoblots demonstrating knockout efficiency of the indicated target at four days post-electroporation are shown below from one representative donor.

**Figure 4.**
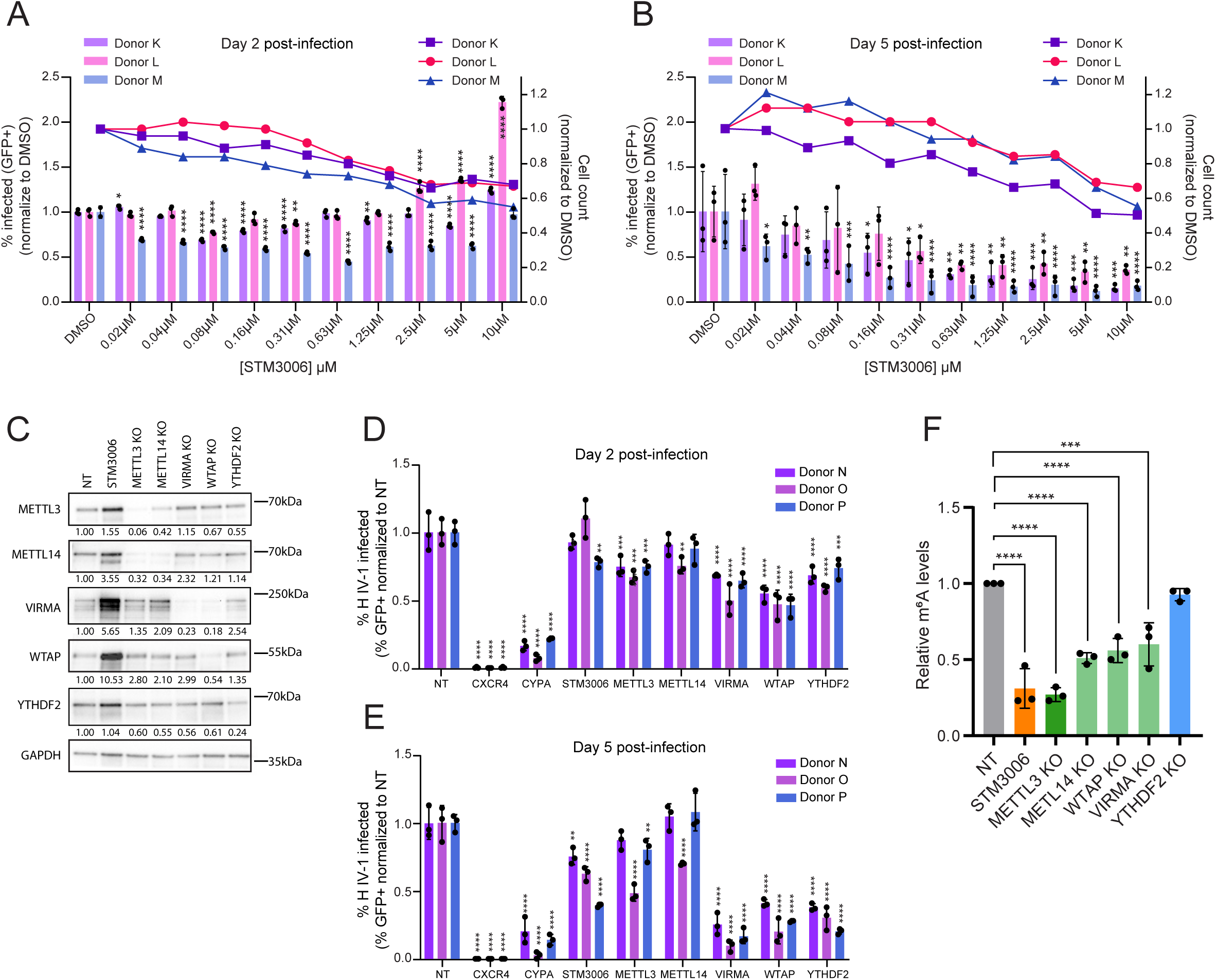
Chemical inhibition, but not knockout of METTL3, reduces HIV-1 infection in primary CD4+ T cells. **A)** Effect of METTL3 inhibitor treatment on viability and HIV-1 infection. Left y axis shows percent HIV-1 infected primary CD4+ T cells at two and **B)** five days post-infection (dpi) in the presence of the indicated concentration range of the METTL3 inhibitor STM3006 in three independent, HIV-1 seronegative donors, measured as percent GFP positive cells by flow cytometry. Cells were pre-treated with STM3006 or DMSO for 24 hours prior to infection. Data represent the mean +/- standard deviation of technical triplicates; statistics were calculated relative to the NT control per condition by one-way ANOVA with Dunnet’s test for multiple comparisons. Right y-axis shows the cell of the same samples, shown by cell count on the same day as the infection data, measured by flow cytometry, normalized to DMSO vehicle control treated cells. **C)** Representative immunoblot of primary CD4+ T cells from one donor, demonstrating protein levels of the indicated m^6^A regulatory protein targets at four days post-electroporation with the indicated CRISPR-Cas9 gene editing crRNPs, or cells treated with 500nM STM3006 for 24 hours. Values below each band show the quantification, normalized to the GAPDH loading control and the NT condition. **D)** Comparison of METTL3 inhibitor treatment and knockout effect on HIV-1 infection. Percent HIV-1 infected primary CD4+ T cells at two and **E)** five dpi in the presence of the indicated perturbation in three independent, HIV-1 seronegative donors. Cells were infected four days after electroporation for knockout conditions, and 24 hours after STM3006 pre-treatment. Data represent the mean +/- standard deviation of technical triplicates; statistics were calculated relative to the NT control per condition by one-way ANOVA with Tukey’s test for multiple comparisons, comparing all conditions. **F)** Total m^6^A levels in poly(A) enriched RNA as measured by m^6^A ELISA in the indicated knockout and STM3006 treated cells at five dpi with HIV-1. Data represent the mean +/- standard deviation of biological triplicates (n = 3 donors); statistics were calculated relative to the NT control per condition by one-way ANOVA with Dunnet’s test for multiple comparisons.

### Data and code availability

No omics datasets were generated by this project. Full results of the CRISPR-Cas9 screen can be found in Table S2. No software was generated for this project. This paper does not report original code. Subfigures and plots were generated using GraphPad Prism 10 (GraphPad Software, La Jolla, CA). Diagrams were generated using BioRender.com. All figures were assembled in Adobe Illustrator (Adobe, Inc.).

### Ethics statement

PBMCs were obtained from anonymous, deidentified blood donors purchased from Vitalant or STEMCELL Technologies.

## RESULTS

### mRNA of the core m^6^A regulatory genes are expressed in primary CD4+ T cells

Previous studies on the role of the m^6^A machinery in HIV-1 replication in diverse cell models have drawn contradictory conclusions, likely due to m^6^A regulation being heavily dependent on the cellular context (33). Therefore, we first wanted to characterize the m^6^A regulatory landscape by determining the mRNA expression level of members of the core m^6^A machinery (**Figure 1A**) in untreated, activated, and type I interferon (IFN)-stimulated primary CD4+ T cells. CD4+ T cells were isolated from peripheral blood mononuclear cells (PBMCs) from two independent HIV-1 seronegative donors. A portion of these cells were activated with plate-bound anti-CD3 antibodies and soluble anti-CD28 antibodies for 72 hours. A portion of these activated CD4+ T cells were then additionally stimulated with type I IFN for 24 hours. RT-qPCR was used to quantify mRNA levels of the core m^6^A machinery relative to the housekeeping gene *TBP*.

We first assessed the basal mRNA expression levels. Of the 14 genes tested (four methyltransferases, two writer complex auxiliary members, one writer inhibitor, two erasers, and five readers), all were found to be expressed, and at comparable mRNA levels in both donors (**Figure 1B**). Amongst the writers *METTL14* had the highest mRNA expression and was roughly 3-fold more abundant than *METTL3*. *METTL16* was found at similar levels to *METTL3*, whilst the small nuclear RNA (snRNA) methyltransferase *METTL4* had the lowest mRNA expression. Between the two erasers, *ALK5BH* was expressed roughly 5-fold more highly than *FTO*. Amongst the core YTH family of readers, *YTHDC1* had the highest expression, followed closely by *YTHDF1*, whilst *YTHDC2* had the lowest mRNA expression (**Figure 1B**).

To investigate the effect of T cell activation and exogenous IFN on the m^6^A machinery, the comparative Ct method was used to calculate gene expression fold changes relative to the untreated condition in each donor’s cells (**Figure 1C**). The IFN-stimulated genes (ISGs) *MX1* and *OAS1* were used as positive controls for response to IFN stimulation. Both *MX1* and *OAS1* increased upon T cell activation in both donors and further increased upon IFN stimulation as expected. In contrast, the expression of the majority of m^6^A machinery genes did not significantly change upon T cell activation or IFN stimulation. The sole exception was the eraser *FTO* which significantly increased upon T cell activation in both donors (fold change = 4.52, p=0.0004 and fold change = 5.15, p<0.0001 respectively).

These results indicate that all core members of the m^6^A writer complex, the other related m^6^A methyltransferases, both erasers and the core readers are expressed at the transcript level in primary CD4+ T cells. Furthermore, expression of most of these factors is unaffected by cell state, however T cell activation triggers an increase in *FTO* expression.

### Arrayed CRISPR-Cas9 screen of m^6^A/m^5^C factors regulating HIV-1 replication in primary CD4+ T cells

To understand the effect of m^6^A and m^5^C factors on HIV-1 replication in primary CD4+ T cells, we performed an arrayed CRISPR-Cas9 knockout screen of 46 genes previously suggested to be involved in m^6^A or m^5^C modification of RNA (**Table S2**). Four independent guide RNA (gRNA) per gene were multiplexed and complexed with Cas9 to generate CRISPR-Cas9 ribonuclear proteins (crRNPs). These were delivered to activated, primary CD4+ T cells from four independent, HIV-1 seronegative donors by electroporation (**Figure 2A**), allowing for the targeted ablation of gene expression with limited toxicity and without the need for selection (49). A non-targeting (NT), gRNA that does not align to any PAM-adjacent region of the human genome was included as a negative control and a previously validated gRNA targeting the HIV-1 co-receptor CXCR4 was used as a positive control.

Representative knockout efficiency of several members of the writer complex (METTL3, METL14, WTAP, and VIRMA) was validated four days post-electroporation by immunoblot (**Figure 2B**), revealing both positive and negative co-dependency. Knockout of METTL3 also reduced the steady-state level of METTL14, while knockout of WTAP reduced the steady-state level of METTL14 and VIRMA. Conversely, knockout of either WTAP or VIRMA increased the level of METTL3 and knockout of METTL3 increased the level of WTAP and VIRMA (**Figure 2B**). At four days post-electroporation, cells were infected in technical triplicate with replication-competent fluorescent reporter virus HIV-1 NL4-3 Nef:IRES:GFP (50). Live cell count (to assess viability) and percent GFP+ cells (to measure percent infection) were monitored by flow cytometry at two, five and seven days post-infection (dpi). As shown at two dpi, knockout of most genes had minimal impact on viable cell number across all donors, except the m^6^A reader HNRNPC and the methyltransferase METTL4 in one donor, which were excluded from further analysis (**Figure 2C**).

The log2 fold change of percent infected (GFP+) cells was calculated relative to the median for each plate and averaged across technical replicates for each donor. Each gRNA pool was then rank ordered by its level of effect on HIV-1 infection at two, five and seven dpi (**Figure 2D**). The majority of gRNAs clustered closely around the plate median (dashed red line) including the four NT control gRNAs (black dots). Knockout of *CXCR4* (orange dots) resulted in strong, reproducible decreases in infection across all timepoints, in all donors as expected. The mean log2 fold change for each knockout across the four donors was then calculated with the threshold for candidate hit calling defined as greater than 2 fold from the median, i.e. at least a 100% increase or a 50% decrease in HIV-1 infection (**Figure 2E**). At two dpi, only knockout of the positive control *CXCR4*, met this threshold, suggesting the m^6^A machinery has limited impact on early stages of HIV-1 infection (**Figure 2E**, left panel). At later timepoints, five and seven dpi, three and four genes met this threshold, respectively. The only knockout identified to increase HIV-1 infection using this criteria was the writer METTL16 thought to have limited methyltransferase activity (28–30), which led to an average of 152% increase in HIV-1 infection at five dpi (log2 fold change = 1.34, **Figure 2E**, middle panel), suggesting METTL16 has antiviral activity against HIV-1 in primary CD4+ T cells. The increase in infection was somewhat variable between donors but each donor exhibited the increase only at the five dpi timepoint (**Figure 2F**; **Table S2**), suggesting the effect is somewhat transient. The other hits identified all led to decreases in HIV-1 infection and include the reader YTHDF2 which promotes degradation of m^6^A-containing mRNAs, and three members of the writer complex WTAP, VIRMA and ZC3H13 (**Figure 2E**, middle and right panel). While there was some donor variability regarding the magnitude of the effect (**Figure 2F**; **Table S2**), knockout of all four genes led to a consistent decrease in HIV-1 replication across all four donors, suggesting they act as HIV-1 dependency factors in primary CD4+ T cells. Surprisingly, knockout of METTL3 and METTL14 that make up the core heterodimer writer complex, did not reach the hit criteria in this screen despite high knockout efficiency (**Figure 2B**) and had very minimal effects on HIV-1 infection (**Figure 2F; Table S2**). In addition, despite successful knockout of *NSUN2*, the primary m^5^C writer (**Figure S2A**), neither NSUN2 nor any of the other m^5^C regulators tested were identified as hits in this screen (**Table S2**).

These results suggest that several members of the m^6^A machinery, namely YTHDF2, WTAP, VIRMA, and ZC3H13, act as HIV-1 dependency factors in activated, primary CD4+ T cells at later stages of infection. The methyltransferase METTL16 appears to have antiviral activity, while knockout of the catalytic core writer METTL3-METTL14 heterodimer itself is less impactful on HIV-1 infection.

### Knockout of WTAP, VIRMA or YTHDF2 significantly reduces HIV-1 replication in primary CD4+ T cells

Several prior reports have found that the core m^6^A writer complex is necessary for HIV-1 replication in different cell line models (35–37,51), which runs contrary to the results from our screen. Therefore, to validate the hits from the screen and to further probe the function of the core m^6^A factors, we repeated our knockout experiments in an additional four independent, HIV-1 seronegative donors, this time using a panel of five individual gRNA per gene in array format (**Figure 3A**). *CXCR4*-targeting and NT gRNAs were included as controls, and protein lysates were collected at four days post-electroporation to assess knockout efficiency by immunoblot. Viability effects were measured by viable cell count at two dpi as above. Cell viability was generally maintained across all knockout conditions, though some individual gRNAs reduced viable cell count across in some donors, and were excluded from downstream analysis (*i.e.*, *VIRMA* gRNA #1, *YTHDF2* gRNA #2, etc., **Figure 3B**). Infection with HIV-1 NL4-3 Nef:IRES:GFP was performed in technical triplicate and, five dpi, the log2 fold change in HIV-1 infection (percent GFP+ cells) for each gRNA was calculated relative to the plate median, averaged across technical replicates and donors (**Figure 3C**). Immunoblots validated efficient knockout of protein expression from each target gene by at least one gRNA in each panel, and one representative donor is shown below the infection data for each target (**Figure 3C**).

Knockout of the positive control *CXCR4* resulted in strong decreases in infection across all donors as expected (average log2 fold change = −9.91, p<0.0001, **Figure S3A**). Knockout of the writers *METTL3*, *METTL14*, *METTL16 and ZC3H13* resulted in no significant change in HIV-1 replication relative to the NT control. Similarly, knockout of the erasers, *ALKBH5* and *FTO*, and the regulator *ZNF217* showed no significant impact on HIV-1 replication despite high knockout efficiency by at least one gRNA. On the contrary, knockout of three of the genes identified in our screen (*VIRMA*, *WTAP*, and *YTHDF2*) resulted in reproducible decreases in HIV-1 replication (**Figure 3C**). Knockout of all other genes investigated did not significantly impact HIV-1 infection (**Figure S3B**). The strongest effect on HIV-1 infection was due to knockout of *VIRMA,* an auxiliary member of the m^6^A writer complex that mediates preferential m^6^A mRNA methylation in the 3′UTR and near stop codons (20). *VIRMA*-targeting gRNA #1 resulted in the strongest reduction in protein and led to the largest decrease in infection (log2 fold change = −2.70, p= 0.0093). Knockout of *WTAP*, another writer complex member thought to regulate recruitment of the m^6^A methyltransferase complex to mRNA targets (18,52), had a more muted impact with gRNA #1 resulting in a log2 fold change of −0.74 (p= 0.0030). Knockout of the m^6^A reader *YTHDF2*, an RNA-binding protein that recognizes and destabilizes m^6^A-modified mRNAs (53), likewise caused a significant decrease in HIV-1 replication with gRNA #1 resulting in a log2 fold change of −1.02 (p= 0.0388, **Figure 3C**).

Taken together, these data validate our screen results showing that the m^6^A writer complex auxiliary members VIRMA and WTAP, as well as the reader YTHDF2, facilitate HIV-1 replication in primary CD4+ T cells, whilst the core writer heterodimer of METTL3 and METTL14 is dispensable.

### Chemical inhibition, but not knockout of METTL3 reduces HIV-1 infection in primary CD4+ T cells

METTL3 is the catalytic subunit of the m^6^A writer complex (11,12), and is stabilized through binding to METTL14 (10,11,54). Whereas WTAP and VIRMA are auxiliary components of the m^6^A writer complex that aid or modify METTL3-METTL14 activity (10,17,20,55). Therefore, it was surprising that knockout of *METTL3* and *METTL14* had negligible impacts on HIV-1 replication in this system whilst *WTAP* and *VIRMA* knockouts resulted in significant decreases in infection. This is also contradictory to several reports in cell line models that showed reduced HIV-1 replication upon *METTL3* knockdown (36,37,51). Therefore, we next sought to characterize the impact of METTL3 inhibition on HIV-1 replication in primary CD4+ T cells using the well-described small molecule METTL3 inhibitor STM3006 (56).

We treated activated primary CD4+ T cells from three independent, HIV-1 seronegative donors with a range of concentrations of STM3006 (0.02-10µM), or DMSO vehicle control, in technical triplicate, 24 hours prior to infection with replication-competent HIV-1 Nef:IRES:GFP. Cell viability and percent GFP+ cells (percent HIV-1 infection) were monitored by flow cytometry at two (**Figure 4A**) and five dpi (**Figure 4B**). At two dpi, higher concentrations of STM3006 led to a reduction in cell number compared to DMSO treated HIV-1 infected cells. The highest concentration (10uM) led to a 36.7% average reduction in cell number, and all concentrations of 0.63µM and above led to an average reduction in cell number of at least 20% compared to DMSO treated HIV-1 infected cells (**Figure 4A**). The effect of drug treatment on HIV-1 infection was variable, with a dose-dependent decrease in infection seen in one donor (M) for doses up to 0.63µM (54.9% decrease, p<0.0001) but no clear trend seen with increasing drug concentrations for the other two donors (K and L) (**Figure 4A**).

At five dpi, the effect of the drug treatment on HIV-1 infection was far more robust and reproducible, with a clear dose-dependent reduction in HIV-1 infection evident for all donors (**Figure 4B**). Treatment with 5µM and 10µM STM3006 led to 79.5% and 77.7% average decreases in HIV-1 infection compared to DMSO treated cells respectively, and all concentrations of 0.31µM or higher led to significant decreases in infection in all donors. However, higher doses of drug did lead to cell toxicity, with the highest concentration of 10µM resulting in an average 43.0% reduction in cell number. The highest dose of drug that maintained at least 80% viability across all donors led to an average decrease in infection of 70.5% (**Figure 4B**).

Taken together, these data show that treatment with the METTL3 inhibitor STM3006 results in a clear, dose-dependent inhibition of HIV-1 replication in primary CD4+ T cells, even at levels that retain cell viability. This is in line with published reports of METTL3 inhibition restricting HIV-1 infection or reactivation in cell lines (57,58), but appears contradictory to the minimal effect we observed in METTL3 knockout primary CD4+ T cells.

To directly compare the effects of the METTL3 inhibitor with knockout of METTL3 and other m^6^A machinery, we repeated the CRISPR-Cas9 targeting and treatment with STM3006 in parallel on activated primary CD4+ T cells from an additional three independent, HIV-1 seronegative donors. Based on **Figure 3C**, the most efficient individual gRNA for knockout of *METTL3*, *METTL14*, *VIRMA*, *WTAP* and *YTHDF2* were identified, which we delivered as crRNPs, as above, alongside NT, cyclophilin A (*CYPA*)-targeting, and *CXCR4*-targeting controls. Based on **Figure 4B**, a concentration of 0.5µM STM3006 was chosen, which was the maximum concentration to yield less than 20% loss in viability at five dpi (**Figure S4A**). Effects on protein level were assessed four days post-electroporation of crRNPs or 24 hours post-treatment with STM3006 (**Figure 4C**). All knockouts were highly effective, with strongly diminished protein expression for each target. As before, we observed both positive and negative co-dependency, as knockout of *METTL3* also reduced the level of METTL14 but increased the level of WTAP and VIRMA, and knockout of *WTAP* also decreased VIRMA protein level. Unexpectedly, STM3006 treatment led to an increase in steady-state levels of all m^6^A writers examined, especially VIRMA and WTAP (**Figure 4C**). Cell viability was measured by flow cytometry on the day of infection (**Figure S4B**). Percent GFP+ cells to measure percent infection were monitored by flow cytometry at two (**Figure 4D**) and five dpi (**Figure 4E**). As observed previously, these knockout conditions resulted in minimal viability defects across all donors (**Figure S4B**).

Knockout of the HIV-1 dependency factors CXCR4 or CYPA led to strong decreases in infection at both two dpi (average decreases of 99.7% and 84.9%, both p<0.001 in all donors, **Figure 4D**), and five dpi (average decreases of 99.9% and 87.8%, both p<0.001 in all donors, **Figure 4E**). At two dpi, knockout of *METTL3* had a moderate effect on HIV-1 infection as observed previously (average decrease of 27.9%, p<0.001 in all donors, **Figure 4D**). Knockout of *METTL14* only resulted in a significant change in HIV-1 infection in one donor (24.8% decrease, p=0.0049, **Figure 4D**). STM3006 treatment at this dose likewise only led to a significant reduction in HIV-1 infection in one donor (21.8% decrease, p=0.0015, **Figure 4D)**. In contrast, knockout of *VIRMA*, *WTAP*, and *YTHDF2* led to more substantial, significant decreases in HIV-1 infection by two dpi with average decreases of 39.2% (*VIRMA* knockout, p<0.001 in all donors), 50.5% (*WTAP* knockout, p<0.001 in all donors), and 32.9% (*YTHDF2* knockout, p<0.001 in two donors and p=0.0002 in one donor, **Figure 4D**).

At five dpi, knockout of *METTL3* had a more variable effect on HIV-1 infection across donors with significant decreases in two donors (19.8% decrease, p=0.0029 and 51.8% decrease, p<0.0001 respectively, **Figure 4E**). Knockout of *METTL14* resulted in a significant change in HIV-1 infection only in the same one donor (30.0% decrease, p<0.0001, **Figure 4E**). In contrast, STM3006 treatment led to a significant change in HIV-1 infection in all donors but with a similar variability to *METTL3* knockout (24.6% decrease, p=0.0027, 37.2% decrease p<0.0001 and 61.0% decrease, p<0.0001, respectively, **Figure 4E)**. However, knockout of *VIRMA*, *WTAP*, and *YTHDF2* led to even stronger, significant decreases in HIV-1 infection, reaching a restriction level similar to knockout of the proviral control gene *CYPA*, with average decreases of 82.8% (*VIRMA* knockout, p<0.001 in all donors), 70.5% (*WTAP* knockout, p<0.001 in all donors), and 70.4% (*YTHDF2* knockout p<0.001 in all donors, **Figure 4E**).

Across all our experiments, the impact of *METTL3* knockout on HIV-1 replication has been minimal in comparison to knockout of the writer complex auxiliary factors, *WTAP* and *VIRMA*. These data additionally show a stronger effect on HIV-1 replication by chemical inhibition of METTL3. While the knockout efficiency of METTL3 protein was high in all experiments, it is possible that slow turnover of the protein or of the reversible m^6^A modifications themselves could have minimized the observed effect on HIV-1 replication relative to the other conditions. We therefore wanted to confirm the effect of knockout and inhibition of METTL3 on m^6^A levels in cellular RNA.

We collected RNA from the STM3006 treated and CRISPR edited cells above at five dpi, enriched for mRNA using polyA selection, and quantified the total levels of m^6^A in each RNA pool by ELISA (**Figure 4F**). As expected, STM3006 treatment significantly decreased global mRNA m^6^A levels (68.8% decrease, p<0.0001, **Figure 4F**), confirming that this dose of drug effectively inhibited METTL3 in primary CD4+ T cells. Knockout of *METTL3* was as effective as the inhibitor in reducing m^6^A levels (73.1% decrease, p<0.0001, **Figure 4F**), suggesting that the *METTL3* knockout is effective and there is no redundancy pathway acting to maintain m^6^A levels in these *METTL3* knockout cells. Knockout of METTL14, VIRMA, and WTAP all led to significant decreases in global m^6^A levels of cellular mRNA although to a lesser extent than METTL3 knockout or inhibition (**Figure 4F**). Knockout of the m^6^A reader YTHDF2 had no effect on m^6^A levels as expected.

Taken together, these results show that reduction of global m^6^A levels of cellular mRNA by *METTL3* knockout or inhibition results in only a minimal decrease in HIV-1 replication in primary CD4+ T cells. Knockout of the writer complex auxiliary proteins, WTAP and VIRMA had less effect on global m^6^A levels of cellular mRNA but resulted in a stronger and earlier decrease to HIV-1 infection. This suggests that either VIRMA and WTAP are playing a secondary role as part of another complex that influences HIV-1 replication in primary CD4+ T cells, or that these factors influence the localization of m^6^A modifications and the altered epitranscriptomic landscape upon WTAP or VIRMA knockout drives the decreased HIV-1 replication phenotype.

## DISCUSSION

Understanding how epitranscriptomic modifications are involved HIV-1 replication, persistence, and reactivation is crucial in revealing the ways in which the virus modulates the cell to create a more permissive environment and could help explain how viral latency is achieved and maintained. Unfortunately, previous studies have resulted in dramatically different depictions of the extent, location, and function of m^6^A and m^5^C modification, and their key regulators during HIV-1 infection (35,39–43,59–65). Many of these conflicting results have been attributed to the well characterized cell type and cell state specific functions of m^6^A modification. Here we aimed to clarify the role of the m^6^A machinery during HIV-1 replication using primary CD4+ T cells, the main targets of HIV-1 infection.

We measured the mRNA expression levels of core m^6^A regulatory factors in primary CD4+ T cells and observed expression of all core factors in these cells, although there were significant differences in the level of expression of each factor (**Figure 1A**). The reader proteins YTHDF1, YTHDF2 and YTHDF3 have been suggested to perform redundant functions (66), with YTHDF2 being dominant due to its higher expression level in 293T cells (35) and HeLa cells (67). However, in primary CD4+ T cells we see *YTHDF1* has consistently higher expression, with *YTHDF2* expression comparable to *YTHDF3*. Despite this, knockout of *YTHDF1* or *YTHDF3* did not affect HIV-1 replication in these cells (**Figure S3B**). We also found that *FTO* mRNA expression in resting, primary CD4+ T cells was significantly lower than *ALKBH5*, however, upon T cell activation it was significantly upregulated, suggesting that in primary CD4+ T cells *FTO* expression is controlled in a cell activation dependent manner. This could be due to the effect of Fe2+ as FTO has been confirmed to exhibit a self-regulatory mechanism by binding to its own gene promoter and repressing its transcription and this binding is prevented by Fe2+ (68) which is taken up by T cells within a minute of T cell receptor (TCR) engagement (69). The functional consequences of increased FTO in activated T cells remain to be investigated, however it has been shown that HIV-1 Gag recruits FTO to remove m^6^A from HIV RNA to promote viral genome packaging (39) suggesting low FTO levels in resting CD4+ T cells could add to the barriers to productive HIV infection of these cells.

We also observed a co-dependency of some writer proteins with *METTL3* knockout and to some extent *METTL14* knockout leading to loss of both proteins. Similarly, knockout of *WTAP* also led to loss of VIRMA, although this was not reciprocated. This relationship between WTAP and VIRMA levels has previously been reported by others (70) and suggests a co-stabilization of the proteins. This is likely also the case for METTL3 and METTL14 which are known to be stabilized by binding to each other (10,11). This could mean that the decrease in HIV-1 infection seen in *WTAP* knockout cells could be due to loss of VIRMA and also suggests that a potential METTL3-independent complex likely contains both VIRMA and WTAP. Similarly, treatment of primary CD4+ T cells with the METTL3-specific inhibitor STM3006 led to an increase in the protein levels of all m^6^A writers probed for, including METTL3 and especially WTAP and VIRMA. In line with the co-dependency phenotypes we observed, we propose this is due to a feedback mechanism whereby the inhibitor binding to METTL3 affects the integrity of the whole m^6^A writer complex, leading to decreased stability of several members which triggers a compensatory increased protein production.

To systematically identify RNA modification factors that influence HIV-1 infection, we performed an arrayed CRISPR-Cas9 screen in primary CD4+ T cells from multiple donors of 46 genes implicated in either m^6^A or m^5^C modification in different systems. As expected, knockout of most of the factors in the screen did not have significant impact on HIV-1 replication. This is unsurprising because HIV-1 relies on a relatively small subset of host proteins for its replication cycle, and furthermore many genes are partially or fully redundant, leading to a minimal impact on HIV-1 replication. HIV-1 is also a highly adaptable virus with mechanisms to bypass various host restrictions. Therefore, even when knockout of a relevant host factor is successful, the virus may use alternative pathways or proteins to maintain replication. We found that only the auxiliary writer complex members WTAP and VIRMA, and the m^6^A reader YTHDF2, significantly influenced HIV-1 infection in our screen. Consistent across our knockout and inhibitor experiments, we saw a stronger effect at five days post-infection (dpi), suggesting the m^6^A factors identified are affecting a later step in the HIV-1 replication cycle, in line with regulation of viral transcripts as shown by many groups (35–37,39,40,59).

There are conflicting reports on whether YTHDF2 enhances or limits HIV-1 infection. Our results are in agreement with several previous studies including one which found knockdown of YTHDF2 inhibited, while YTHDF2 overexpression enhanced, HIV-1 RNA and protein expression, and subsequently viral replication (35). A follow up paper from the same group confirmed YTHDF2 overexpression led to increased viral RNA and viral reporter protein expression as early as 24 hours post-infection with replication-competent HIV-1, and showed this was due to YTHDF2 binding to m^6^A sites on HIV-1 transcripts resulting in stabilization of these viral RNAs (40). However our results contradict those of several other reports including one that found YTDF2 bound to HIV-1 genomic RNA is packaged into virions where it can inhibit reverse transcription (37) and a follow up paper from the same group that found YTHDF2 overexpression in HIV-1 target cells decreased viral genomic RNA levels and inhibited both early and late reverse transcription (71). A report from a different group found that knockdown of METTL3, WTAP, VIRMA or YTHDF2 led to increased levels of HIV-1 RNA (70). We propose these inconsistent phenotypes are likely due to differing cell types and approaches used in these studies, however one study that used primary CD4+ T cells did see an opposing YTHDF2 phenotype (37). In that study shRNA was used to knockdown YTHDF2 prior to infection with a single cycle HIV-1 pseudotype and the authors conclude that the YTHDF2 phenotype is due to it blocking viral reverse transcription. Since we used a replication competent HIV-1 it could be that the reverse transcription inhibition is being masked by a stronger phenotype post-integration, especially since they measured infection at 24 hours post-infection and we see the greatest effects at five dpi.

It is surprising and potentially counterintuitive that the auxiliary writer complex proteins WTAP and VIRMA had a stronger effect on HIV-1 infection than the catalytic subunit METTL3 or its heterodimer partner METTL14. The majority of HIV m^6^A studies have shown that knockdown or inhibition of METTL3 has an inhibitory effect on HIV (35–37,51). We originally theorized that *METTL3* knockout may not be leading to reduced global m^6^A levels perhaps through a redundancy mechanism, however this was not the case as *METTL3* knockout cells had significantly reduced global levels of cellular RNA m^6^A and lower levels than *WTAP* or *VIRMA* knockout cells (**Figure 4F**). The METTL3-METTL14 heterodimer has an RNA sequence specificity to GGACU sites (10), but only a small portion of consensus sites across the transcriptome are methylated (20,72). Several studies have identified auxiliary writer complex members as important for localization. For example, VIRMA was shown to recruit the m^6^A writer complex specificity to 3′UTR regions and near stop codons, and interacts with polyadenylation cleavage factors CPSF5 and CPSF6 to facilitate correlation of m^6^A methylation with alternative polyadenylation (20). Furthermore, WTAP has been identified as crucial for localization of the writer complex into nuclear speckles associated with mRNA export, and the dynamic regulation of m^6^A levels in mRNAs that occur there (6,10,18). However, it has been suggested that loss of these localizing members leads to loss of global m^6^A, for example, knockdown of *WTAP* was previously shown to lead to a larger decrease in total m^6^A levels in HeLa and 293FT cells than knockdown of *METTL3* or *METTL14* (10). In this study we observed a significant proportion of global cellular RNA m^6^A still present in *WTAP* and *VIRMA* knockout cells (**Figure 4F**), suggesting that the infection phenotype is not simply due to widespread loss of m^6^A modification. While the writer proteins are often described as a single writer complex, our data strongly indicates that multiple independent complexes can play a role in HIV-1 replication. This suggests potential writer complex redundancy or heterogeneity, that the m^6^A machinery may form distinct regulatory complexes in primary CD4+ T cells with different combinations of the subunits that could lead to m^6^A modification in distinct locations and dictate different phenotypic outcomes.

Alternatively, like a growing list of histone modifiers, several components of the m^6^A writer complex have been found to perform “moonlighting” functions that are methylation independent. For example, METTL3 promotes translation by directly binding the 3′ UTRs of target mRNAs and interacting with the translation initiation machinery, independent of its catalytic activity (73). WTAP was originally characterized as a splicing regulator (74) and is known to influence alternative splicing and RNA stability in ways separate to its function in m^6^A regulation (75–77). VIRMA interacts with polyadenylation factors and may help couple 3′ end processing with transcript fate independently of methylation (20). Furthermore, the homolog of VIRMA and the homolog of WTAP form part of a complex that regulates alternative splicing in *Drosophila melanogaster* (74). Taken together, it is worth considering that the HIV-1 infection phenotype we are seeing is explained by a METTL3-independent function of WTAP and VIRMA, potentially in regulating splicing.

Despite previous studies suggesting key roles for the two m^6^A erasers in HIV-1 infection (36,37,39,40), even though several gRNAs resulted in substantial knockout of the FTO and ALKBH5 proteins, no significant changes in HIV-1 infection were observed (**Figure 3C**), suggesting that m^6^A erasers are of limited importance during HIV-1 infection of primary CD4+ T cells.

Finally, the methyltransferase NSUN2 serves as the primary writer for m^5^C on HIV-1 RNAs (42,78) and previous studies have shown that NSUN2 inactivation inhibits m^5^C addition to HIV-1 transcripts and also viral replication, via reduced HIV-1 protein levels, which in turn correlates with reduced ribosome binding to viral mRNAs (42). However, despite efficient knockout of *NSUN2* (**Figure S2A**), we did not find an effect on HIV-1 infection in primary CD4+ T cells. Additionally, no other m^5^C factors that were tested had an impact on HIV-1 infection, including the m^5^C RNA methyltransferase NSUN1 which has also been identified as an HIV-1 restriction factor, although in the context of viral latency (43). Like m^6^A, the effect of m^5^C methylation has repeatedly been shown to be cell type specific which may explain the discrepancy between our findings and the literature.

As with all high throughput techniques, it is likely that subtle effects on HIV-1 infection could be missed due to lack of significance, especially for cellular genes with modest or context-dependent roles. This is clearly shown by the modest effect of *METTL3* knockout seen in individual experiments that was not significant in the screen. Furthermore, this study focused on the host RNA methylation factors that influence HIV-1 infection at early stages. It is possible that some factors that had no effect on HIV-1 productive infection in our screen do influence HIV-1 latency or reactivation (57), and further investigation is needed to determine the role of m^6^A in HIV-1 latency and reactivation.

Overall, our results clarify the role of epitranscriptomic machinery during HIV-1 replication of primary CD4+ T cells and suggest that regulation by auxiliary members of the m^6^A writer complex is more influential than the function of the catalytic core itself on HIV-1 infection in primary CD4+ T cells.

## AUTHOR CONTRIBUTIONS

Conceptualization, J.F.H.; Data Curation: K.A.J.-J.; Formal Analysis: K.A.J.-J., L.M.S.; Funding Acquisition: J.F.H.; Investigation: K.A.J.-J., L.M.S., S.H., T.L.J., A.M.S.; Methodology: K.A.J.-J., L.M.S., S.H.; Project Administration: J.F.H.; Resources: J.F.H.; Supervision: J.F.H., L.W.; Validation: K.A.J.-J., L.M.S.; Visualization: K.A.J.-J.; Writing - Original Draft Preparation: K.A.J.-J., J.F.H.; Writing - Review & Editing: K.A.J.-J., L.M.S., L.W., J.F.H.

## ACKNOWLEDGMENTS

We would like to thank Stacia Phillips for advice regarding this work. This research was supported by NIH/NIAID funding for the HIV Accessory & Regulatory Complexes (HARC) Center (U54 AI170792, J.F.H.), NIH funding for the Third Coast Center for AIDS Research (P30 AI117943, J.F.H.), NIH/NIAID grants for HIV research (R01 AI167778, R01 AI150455, R01 AI165236, and R01 AI150998, J.F.H.), and NIH grants (R33 AI169659 and P30 CA086862-25S1, L.W.).

---

**Figure S1** Flow cytometry gating strategy Flow cytometry data were exported as FCS3.0 files using Attune NxT Software v3.2.0 and analyzed with FlowJo™ Software (BD Biosciences) using this gating strategy. **A)** Cells were gated to exclude cell debris by light scatter and the number of cells recorded for viability measurement of each condition. Cells from gate 1 were then filtered for doublet discrimination in both **B)** forward and **C)** side scatter. **D)** Cells with equal fluorescence in the BL-1 (GFP) channel and the VL-1 channel were identified as autofluorescent and excluded from the analysis. A consistent gate was then used to quantify the fraction of remaining cells that expressed GFP.

**Figure S2** Validation of knockout of the m^5^C writer NSUN2 **A)** Western blot showing knockout efficiency of the m^5^C writer *NSUN2* in primary CD4+ T cells from four independent, HIV-1 seronegative donors, measured four days post-electroporation with a multiplexed pool of five individual CRISPR-Cas9 ribonuclear proteins (crRNPs) targeting *NSUN2* (N2) or the control gene *CXCR4* (X4).

**Figure S3** Validation of CRISPR-Cas9 screen using independent gRNAs **A)** HIV-1 infection of primary CD4+ T cells from four independent, HIV-1 seronegative donors, measured two days post-infection (dpi) with replication competent HIV-1 NL4.3, four days post-electroporation of CRISPR-Cas9 ribonuclear proteins (crRNPs) targeting the HIV-1 dependency factor and positive control *CXCR4*, shown as % infected (GFP+) cells normalized to the non-targeting (NT) control condition, measured by flow cytometry. Each dot represents the log2 fold change in infection for one donor, bar height shows average of four donors and error bars show standard deviation. **B)** HIV-1 infection of primary CD4+ T cells from four independent, HIV-1 seronegative donors, measured five dpi with replication competent HIV-1 NL4.3, four days post-electroporation of 3-5 individual crRNPs targeting the indicated genes, shown as % infected (GFP+) cells normalized to the NT control condition, measured by flow cytometry. Each dot represents the log2 fold change in infection for one donor, bar height shows average of four donors and error bars show standard deviation.

**Figure S4** Comparison of viability of knockout cells to METTL3 inhibitor treated cells **A)** Cell viability of primary CD4+ T cells from three independent, HIV-1 seronegative donors, measured five days post-infection (dpi) with replication competent HIV-1 NL4.3, 24 hours post-treatment with the METTL3 inhibitor STM3006 at the indicated doses, shown as cell count normalized to DMSO condition, measured by flow cytometry. Dotted lines indicate 80% cell viability (y=0.8) and the highest concentration of STM3006 to yield at least 80% cell viability in all donors (x=0.44µM). **B)** Cell viability of primary CD4+ T cells from three HIV-1 seronegative donors, measured four days post-electroporation of CRISPR-Cas9 ribonuclear proteins (crRNPs) targeting the indicated genes or 24 hours post-treatment with the METTL3 inhibitor STM3006 at 0.5 µM, shown as percent viable cells, measured by flow cytometry. Each dot represents one donor, and lines show average of three donors and error bars show standard deviation.

**Table S1 List of Antibodies, Primers and crRNAs**

**Table S2 Results of Multiplexed CRISPR-Cas9 Screen**

