## Supplementary figures and images for "Impact of N^6^-methyladenosine (m^6^A) machinery on HIV-1 replication in primary CD4+ T cells"

### Supplemental Figures

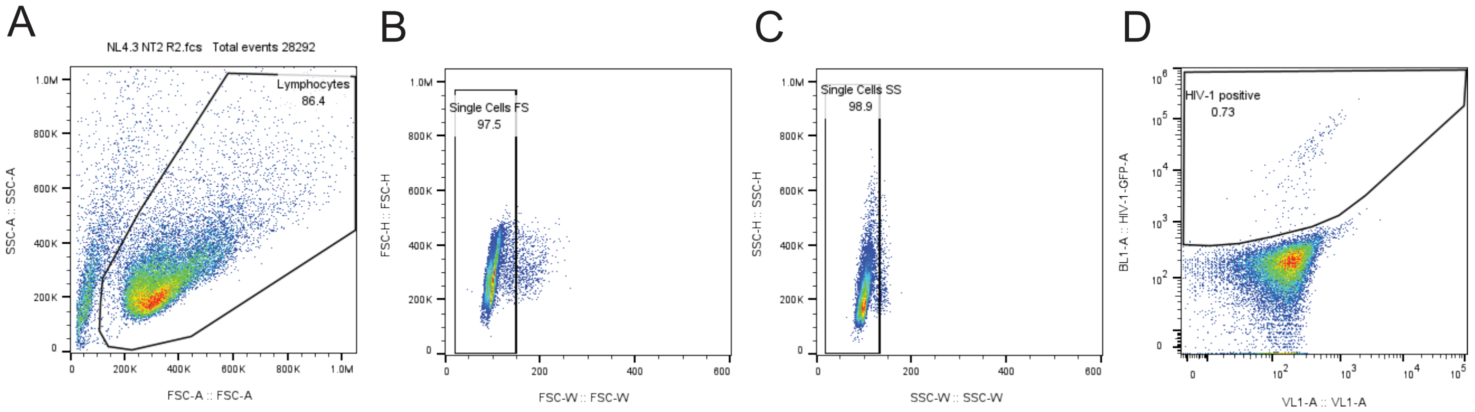

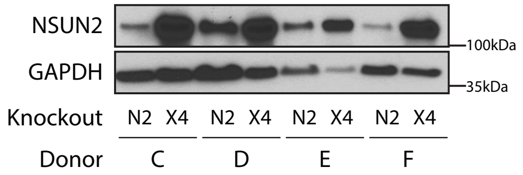

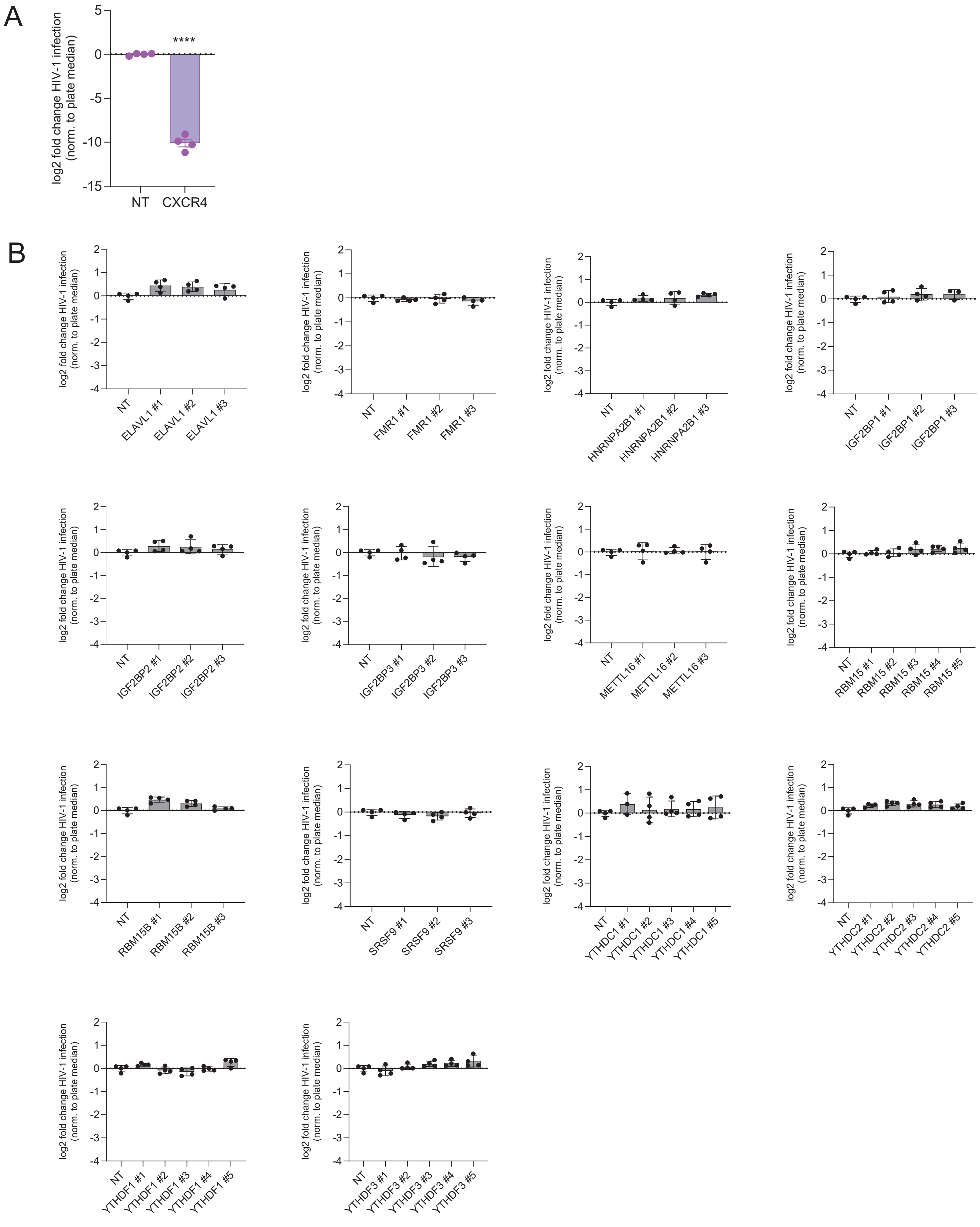

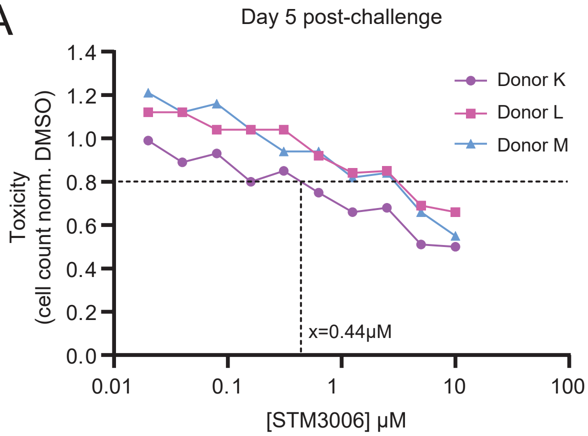

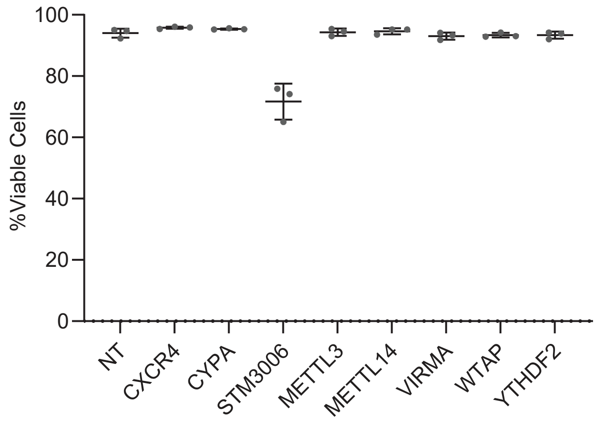
